# SpikeScape: A Tool for Analyzing Structural Diversity in Experimental Structures of the SARS-CoV-2 Spike Glycoprotein

**DOI:** 10.1101/2022.11.15.516662

**Authors:** Darya Stepanenko, Carlos Simmerling

**Affiliations:** Laufer Center for Physical and Quantitative Biology; Department of Applied Mathematics and Statistics, Stony Brook University; Department of Chemistry, Stony Brook University

## Abstract

In this application note we describe a tool which we developed to help structural biologists who study the SARS-CoV-2 spike glycoprotein. There are more than 500 structures of this protein available in the Protein Data Bank. These structures are available in different flavors: wild type spike, different variants, 2P substitutions, structures with bound antibodies, structures with Receptor Binding Domains in closed or open conformation, etc. Understanding differences between these structures could provide insight to how the spike structure changes in different variants or upon interaction with different molecules such as receptors or antibodies. However, inconsistencies among deposited structures, such as different chain or sequence numbering, hamper a straightforward comparison of all structures. The tool described in this note fixes those inconsistencies and calculates the distribution of the requested distance between any two atoms across all SARS-CoV-2 spike structures available in the Protein Data Bank, with the option to filter by various selections. The tool provides a histogram and cumulative frequency of the calculated distribution, as the ability to download the results and corresponding PDB IDs.

## Introduction

The urgency for COVID-19 treatment and vaccine development led scientists to determine many cryo-EM structures of the SARS-CoV-2 spike protein during a relatively short period of time. Insight into details of protein structure, and mechanisms of protein action, can be highly valuable for target-based drug discovery, vaccine development, exploring differences between variants to predict vaccine effectiveness, and for understanding viral infection mechanisms in general. As of October 19th, 2022 there were 585 structures of the SARS-CoV-2 spike protein in the Protein Data Bank (PDB) with a resolution better or equal to 4.0Å and mass greater than or equal to 400*kDa*. Those spike protein structures are present in different flavors including wild type spike, 2P mutants, different variants, structures with bound antibodies, structures with Receptor Binding Domain in closed or open conformation, etc.

To identify similarities and spot differences between all structures, scientists typically need to load structures into molecular visualisation software such as VMD,^1^ Pymol^2^ or Chimera^3^ for visual inspection. The spike protein is a large glycoprotein as shown in Figure 1; it consists of 3 chains, each with 1273 amino acids.^4,5^ Visually comparing pairs of very large structures might be tractable, but comparing 585 structures becomes daunting. Moreover, all full-spike structures currently available in the PDB are cryo-EM models, with not very high resolution. This means that differences noted between pairs of structures might be significant, or might simply reflect structure uncertainties. In addition, the resolution is not uniform across each structure, and there are some regions that are less reliable than others in different structures.

**Figure 1:**
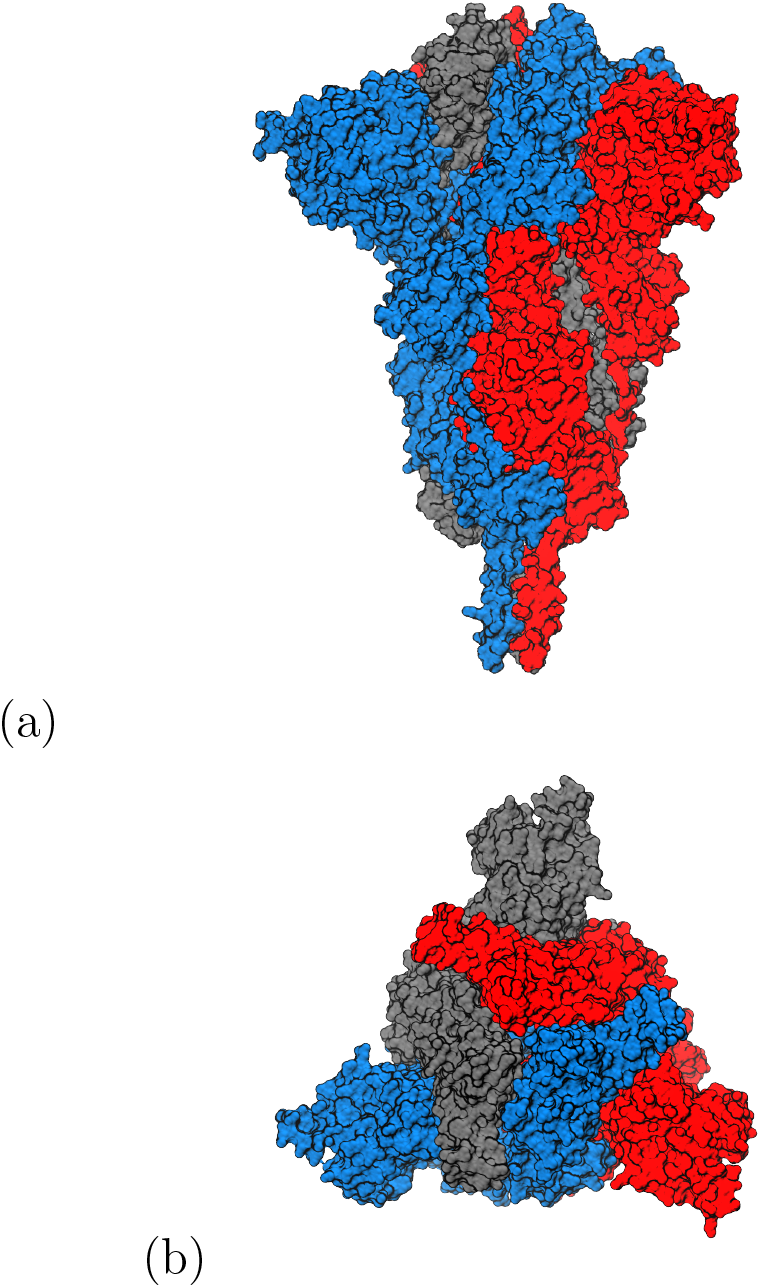
Surface representation of the SARS-CoV-2 spike glycoprotein trimer colored by protein chain: (a) side, (b) top views.

It would be helpful to have a tool that could analyze many SARS-COV-2 spike structures simultaneously to identify differences and similarities, separate them by features, and decrease the influence of low-resolution modeling. The tool described here attempts to address this problem. Our tool measures the distance between any two selected atoms in all SARS-CoV-2 spike structures available in the PDB at runtime, and returns a distance distribution to the user. For this to work properly, all spike structures from the PDB should be consistent with each other: sequence numbering, chain identifiers, and chain order in the trimer should match. Below we describe how the tool eliminates chain naming and chain order inconsistency in the program, identifies sequence numbering problems, removes structures where the requested atoms are missing, and provides a consistent analysis of the available structures.

The presented tool can be also used as a filter to create a subset containing only the structures with a specific feature that cannot be filtered by the PDB itself. Examples include filtering only structures with an open Receptor Binding Domain, or with a resolved Receptor Binding Motif, etc. The tool can also filter structures based on the presence of a specific mutation.

## Implementation

### Selecting structures

The first task of the program is to obtain from the PDB a list of available structures using the SARS-CoV-2 spike UNIPROT ID (P0DTC2), mass larger than 400*kDa*, and a resolution better than 4.0Å. The program updates the list of available structures during every run. As we are interested in pre-fusion spike structures only, and there is only one post fusion spike structure in PDB with pdb id 6XRA, the program removes this PDB id from further analysis. Next, the program downloads any PDB files that are not already stored, for example if a new structure appeared since the previous run of the program. The program attempts to download the file in PDB format; if unavailable, the program downloads the mmcif file instead.

### Chains and sequence numbering consistency

The spike structure has 3 chains, as shown in Figure 1. However some structures contain additional proteins such as receptors or antibodies. In such cases, the PDB file has more than 3 chains. An approach is needed to extract the 3 chains of the spike protein. We employ the accepted and widely used spike amino acid sequence numbering. It is also known that coronavirus spike proteins contain highly-conserved disulfide bonds, one of which is C1126-C1082 in SARS-CoV-2. The program uses the known sequence and disulfide information for chain detection. C1126 should be present in each of three chains in any valid spike protein, and we assume that C1126 will be absent in any antibody or receptor chain. The program checks each chain in the PDB file for the amino acid 1126 being a cysteine. Additionally, the tool checks to ensure that amino acid 57 is proline. All chains that have C1126 and P57 are considered to be spike chains. If a structure does not have 3 chains with C1126 and P57 it means that sequence numbering in at least one of the chains is inconsistent with other structures and this structure is removed from further analysis.

In our tests, the program detected 66 structures with the residue 1126 not being cysteine or residue 57 not being a proline in 3 chains. These structures are removed from further analysis. The updated list of removed structures is shown to the user every time the program runs. This list updated on October 19th is shown in the List S1.

### Distances between atoms in the same or different chains

Since the spike protein is a trimer, the distance between a pair of atoms could be calculated either between two atoms within the same chain, or between two atoms from two different chains. These cases are described separately.

### Algorithm for calculating the intra-chain distance

If both atoms of interest are from amino acids in the same chain, the program attempts to measure the distance between selected atoms in each of 3 chains. If the requested atoms are present in all 3 chains, the program will return 3 distances. If a chain does not have at least one of requested atoms, the program will not be able to calculate the distance in this chain and will notify the user, and return distances for the other chains if possible. After the distance is calculated across all structures, the histogram and cumulative histogram are plotted, and a table with all calculated distances and their corresponding PDB IDs is presented.

### Algorithm for calculating the inter-chain distance

#### Chain order assignment

If atoms of interest are in different chains, the order of chains in the trimer needs to be consistent among the structures. It is apparent when viewing the spike from the top, as shown in Figure 1(b), that the structure can have a clockwise or a counter-clockwise order of the three chains, depending on how they are ordered in the PDB file. This order is inconsistent across current PDB files, even when using the chain letters present in the PDB file (chains A-B-C are observed to occur in clockwise order in some PDB files, and counter-clockwise in others, and some structures have chains names different from A, B and C). If the variable order is not taken into account, the distance measurement in one structure could be performed clockwise, while in the other structure it might be performed counter-clockwise. This will make the results in the histogram inconsistent and hamper the desired comparisons.

For that reason we implemented an additional step of assigning chain order in each structure prior to the distance calculation. The tool selects the first chain in the structure, naming it chain A. It then measures the distances between the C*α* at position 971 in chain A to the C*α* at position 752 in each of the other two chains, as shown in Figure 2. The chain for which the distance is smaller is named chain B, and the other chain is named chain C. By measuring these 2 distances and assigning chain order for every structure, the program makes sure that chain ordering is consistent across all structures.

#### Inter-chain distance calculation

After the tool assigns chain order, it selects the first requested atom, in each of the three chains. For each of these, it identifies the other requested atom, in different chains using both the clockwise and counter-clockwise order. Thus there are 6 possible distances for each inter-chain request: 3 clockwise and 3 counter-clockwise. If either of the requested atoms is not resolved, the distance for this pair will not be calculated and the program will notify the user. The program will calculate one histogram for distances calculated in the clockwise order, and another histogram for distances in the counter-clockwise order. The user will determine whether the clockwise or counterclockwise order is most relevant, depending on the scientific question.

**Figure 2:**
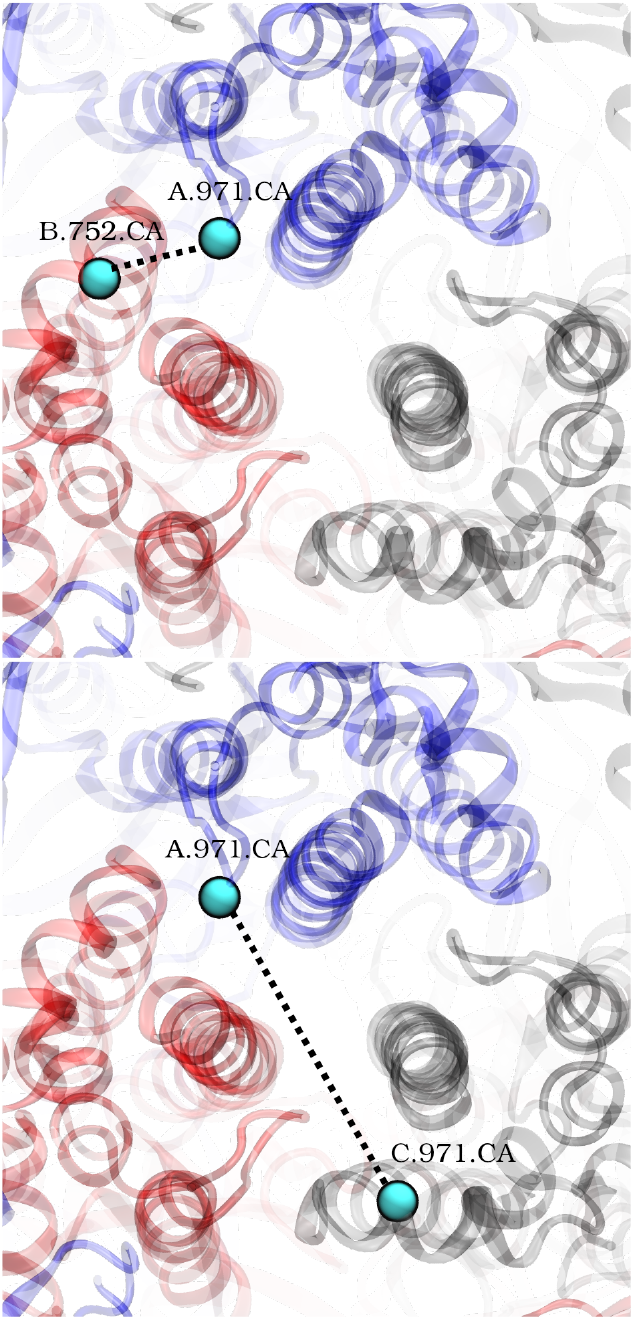
Distance calculation to set up consistent chain order. Each chain has a different color, consistent with Figure 1. A dashed line is drawn between the C*α* atom in amino acid 971 of chain A to C*α* atom in amino acid 752 of the other two chains. Chain identifier B is assigned to the chain with the shorter distance to chain A (top).

### Sequence mutations and substitutions

If a user is interested only in structures with a specific sequence alteration present, for example from a specific variant or with/without a specific experimental modification, there is an option to turn on the mutation filter. The user specifies the amino acid number and name, and the list of analyzed structures that match the requested filter will be presented to the user. Only these structures will be used for the subsequent distance calculation. In this manner one can narrow the analysis to only structures that contain a specific amino acid, which could be either a mutation or the original wild-type sequence.

### Data output

For each distance request, the user will receive separate histogram and cumulative histogram plots for both orders, together with two tables with all calculated distances and the corresponding PDB IDs. These can be downloaded for more detailed analyses. The ranges for the X and Y axes are set automatically to include data from all structures analyzed.

## Examples

A few examples below show how the program can be used to explore spike features. The first example demonstrates how to choose between two histograms when measuring the distance between atoms from different chains. The next two examples show how the tool can identify different conformations, and how they might be influenced by sequence substitutions. The fourth example shows how to use the tool to filter structures by resolved or non-resolved regions. The last example illustrates how the tool checks if an unexpected feature observed in one structure is reproducible in others.

### Example 1: choosing between clockwise and counter-clockwise histograms

This example demonstrates how to interpret the two histograms that are provided when the user requests a inter-chain distance. As discussed above, there are 2 ways to calculate an inter-chain distance as shown in Figure 2: counter-clockwise or clockwise.

The user might want to explore how the N-terminal Domain (NTD) from one chain interacts with the C-terminal Domain-1 (CTD1) from the clockwise chain. In PDB 6XR8,^6^ a hydrogen bond is formed between F43.N from the NTD and F565.O from CTD1 of the clockwise chain. The distance in one order is 2.8 Å, while the distance in the other order is 53.7 Å (Figure 3a). The program will calculate the inter-chain distances for all PDB files, and present separate data for each order. The user can determine which is most appropriate to their query.

By measuring the inter-chain distance F565:O - F43:N, one can investigate the frequency at which this hydrogen bond is present, across all structures. The tool calculates the distances in both chain orders (Table S1, S2) and outputs two distributions as shown in Figure 3. Only one of 2 distributions shown in Figure 3 results in distances consistent with a hydrogen bond (around 2-3ÅA), while the other distribution is spread between 58-65AÅ. In this case the user should identify that the distribution in Figure 3(c) is the relevant one; it confirms that the majority of structures have this hydrogen bond. A few individual structures are outliers with distances longer than 5ÅA, reinforcing the value of measuring the entire distribution.

### Example 2: identifying different conformations

The dynamic motion of the Receptor Binding Domain (RBD)^7–9^ has been widely studied. A user may want to explore the distribution of RBD position across all structures. In the closed state, the RBD of one chain packs against the Central Helix (CH) of the clockwise chain; these move apart when the RBD is open (indicated using PDB 6VYB^7^ in Figure 4(a). To quantify how many structures with open or closed RBDs are present in the PDB, the user could measure the distance from the top of the CH using atom 987.C*α* in one chain, to RBD atom 413.C*α*, in the other chains.

**Figure 3:**
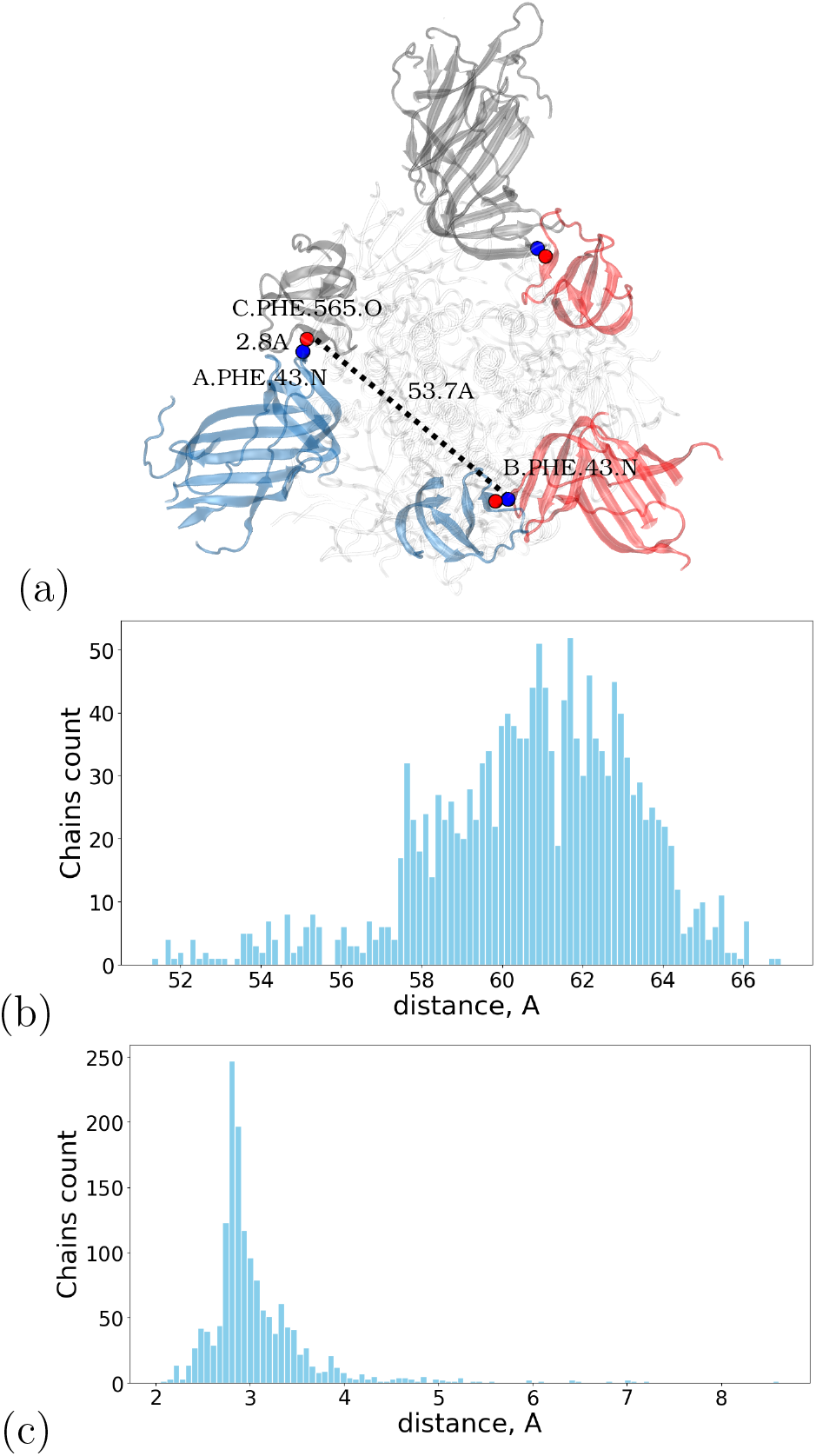
Inter-chain distance between F565.O (red circle) from chain A and F43.N (blue circle) from other chains to investigate the presence of a hydrogen bond. Distances are measured to both other chains (dashed lines). **(a)** Spike structure top view with NTD and CTD1 colored by chain. **(b)** Distribution of the F565.O-F43.N distance in clockwise and **(c)** counterclockwise orders.

As usual, the tool produces two histograms; the one showing a population of short contact distances is selected (Figure 4(a)). Two peaks are apparent, the first one with the maximum at 6AÅ, and the second is spread around 39ÅA. These are consistent with distances in PDB 6VYB of 5.9A for the closed RBD and 36.5Å for the open RBD (Figure 4(a)). The peak corresponding to the open RBD is wider, which illustrates the flexibility of the RBD openness as was originally reported for the SARS-CoV spike.^10^ The table produced by the tool with up to three distances per PDB is provided in Table S3. Using the data in this table, a user can analyze how many RBDs out of 3 are open or closed based on the distance, as well as identify structures with the desired RBD position that could be subjected to additional analyses by the user.

**Figure 4:**
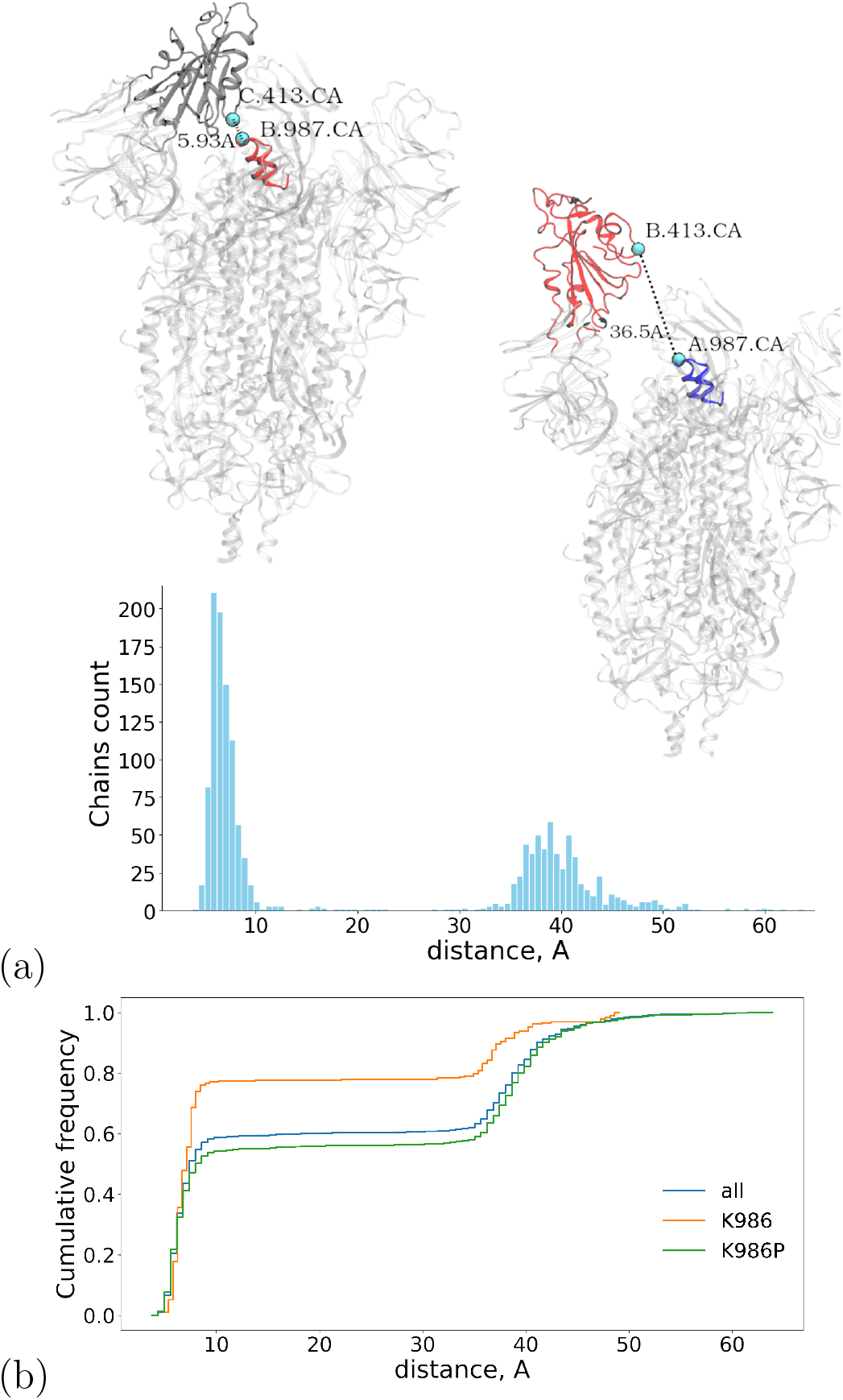
**(a)** (Upper) spike structure (PDB id 6VYB^7^) with closed (on the left) and open (on the right) RBD, color-coded by chain. Non-RBD domains are shown in gray. (Lower) The distance distribution between 987.C*α* and 413.C*α* for all PDB structures shows 2 peaks: the first near 6Å, and the second near 39Å. **(b)** Exploring the impact of sequence variation on dynamics: cumulative frequency of the distance 987.C*α*-413.C*α* in all (blue line), only K986P (green line), and only K986 (orange line) spike structures.

### Example 3: studying structures with a specific mutation

Many SARS-CoV-2 spike proteins used in research and vaccine development have two proline (2P) substitutions (K986P and V987P) that stabilize the prefusion spike structure.^9,11^ It has been suggested that the change in electrostatic interactions from K986P could alter the stability of the closed RBD.^6^ To explore this issue, the measurement explained in the previous example can be performed independently on structures with either proline or (wild-type) lysine at position 986. Such filtering is achieved by entering the desired amino acid name and number before calculating the distance distribution and cumulative histogram.

The tool detected 422 structures with K986P and 97 structures with K986 listed in List S4 and List S6 respectively. Interestingly, these structures subsets exhibit different RBD opening distributions as quantified by the 987.C*α*-413.C*α* distance (Figure 4(b), Table S4, S5). When all structures are included, around 60% of chains are closed. When K986P is present, a little less than 60% of chains are closed. However, with the wild-type amino acid (K986), around 80% of chains adopt a closed RBD. One interpretation of this result is that the 2P substitution leads to increased RBD opening, however there could be other experimental reasons for these differences.

### Example 4: filtering structures by resolved regions

If one is interested only in structures that have a particular region resolved, they can choose one amino acid from this region and check which structures have coordinates for an atom from this amino acid. For example, the coordinates of the first few amino acids in the N-teminal Domain (NTD) are not modeled in many SARS-CoV-2 spike structures. The user may wish to filter only structures with coordinates in this region; here we will use the highly conserved cysteine at position 15. By calculating the distance between C15.C*α* and itself, the tool will output a list of PDB codes with a distance of 0Å for those structures that have coordinates for this atom, and a separate list of structures in which the atom is not present. Using this example, there are 880 chains (in 298 structures listed in List S7) without coordinates for C15.C*α*, and only 228 structures with C15.C*α* being resolved in at least one chain (Table S6).

### Example 5: confirming feature reproducibility

As mentioned above, spike structures are obtained using cryo-EM experiments, and can have limitations based on low resolution in some regions. This can lead to uncertainty about the reliability of an individual structure. If an unexpected feature is observed, the tool can be used to check whether this feature is reproducible in other PDB structures. This analysis can provide more confidence in the individual model, or perhaps suggest that the structure represents either a modeling error, or perhaps an interesting outlier.

We demonstrate this function by examining two cysteine amino acids at positions 1043 and 1032 in the structure 7E5R^12^ (Figure 5(a)). Based on the distance between the sulfur atoms, one might conclude that these cysteines do not form a disulfide bond. The tool can be used to check if the observation is reproducible in other available structures by measuring the distance between SG atoms of C1032 and C1043 in all available PDB structures (Table S7).

As seen from the distance distribution in Figure 5(c,d) nearly all structures adopt a distance of 2.0Å, which corresponds to the disulfide bond as shown in Figure 5(b). However, there are 5 structures in the distribution that have distances higher than 3Å, including 7E5R^12^ (3.9Å, 4.0A, 4.0Å). Other PDB IDs that appear to be outliers for all three disulfide bonds are 7LJR^13^ 7FJN^14^ and 7FJO,^14^ while PDB ID 7U0P^15^ lacks one disulfide bond. All structures except these five show all three disulfide bonds, suggesting that the unusual observation from PDB 7E5R^12^ is poorly reproducible. This might be a a scientifically important observation or a modeling artifact.

**Figure 5:**
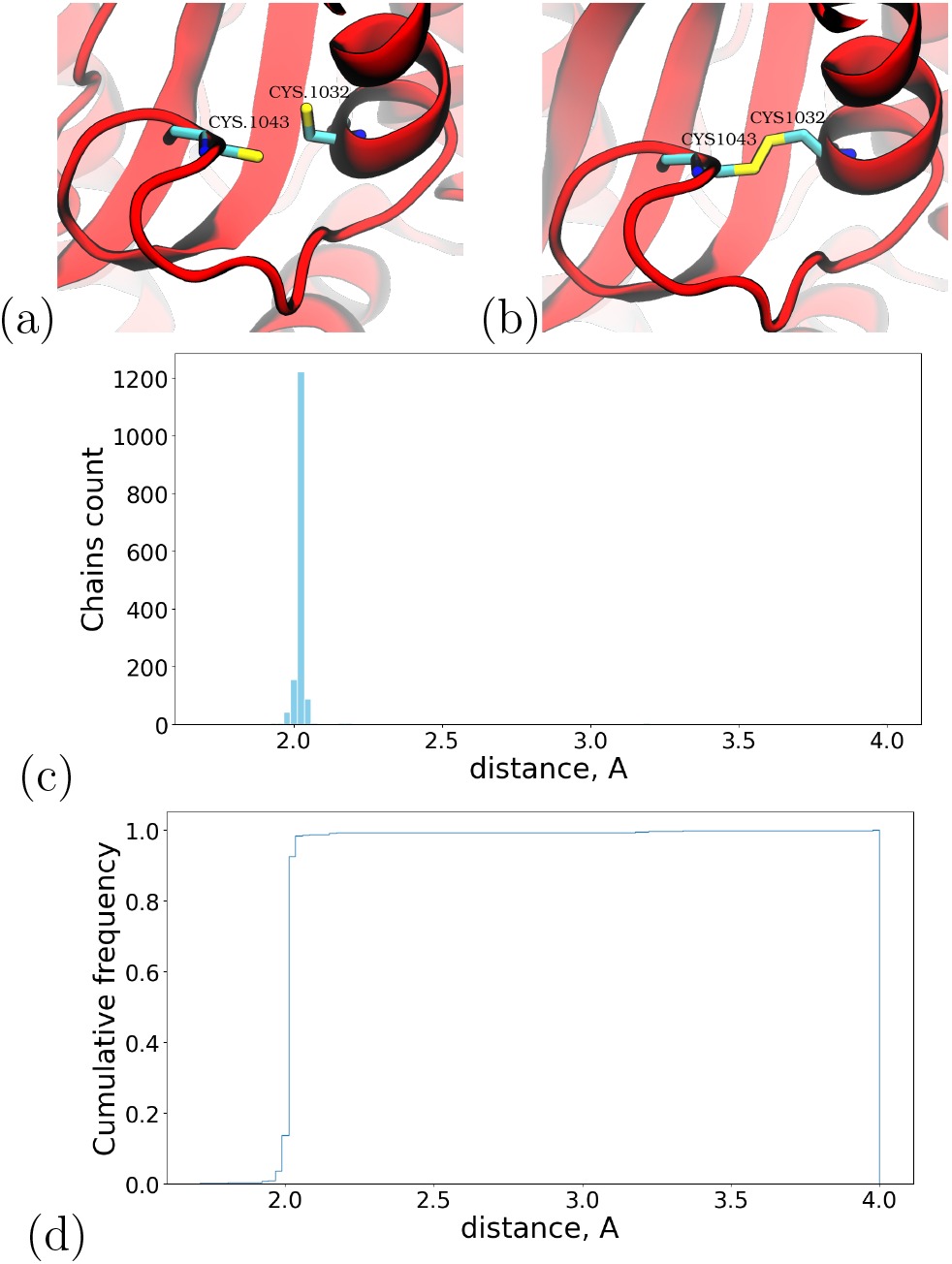
Distance between C1032.SG and C1043.SG. (a) atoms are too far apart to form a disulfide bond (PDB ID 7E5R^12^), (b) atoms form disulfide bond (PDB ID 6VYB
^7^). **(c)** Histogram and **(d)** cumulative frequency of this distance across all structures.

## Conclusion

Experimentalists have produced almost 600 structures of the SARS-Cov-2 spike protein during the COVID-19 pandemic. This large database of structures for one protein allows exploration of how the structure of a protein can vary depending on sequence or experimental conditions, and potentially provide insight into how environment influences the viral infection mechanism. An ability to identify differences and similarities among many structures is crucial for vaccine development as well as variants study. This tool allows rapid comparison of all spike structures with each other, identifying structures with specific features or specific mutations, and exploring the reproducibility of features or interactions. At present the tool measures only the distance between selected atoms but can be extended to other metrics such as angles and dihedrals, or other data present in the PDB files. The approach also could be extended to other proteins, especially those that are widely represented in the Protein Data Bank.

## Supporting information

Supplemental Lists

TableS1.distance_43n_565o_diff1

TableS2.distance_43n_565o_diff2

TableS3.distance_987ca_413ca_diff1

TableS4.distance_987ca_413caLYS986_diff1

TableS5.distance_987ca_413caPRO986_diff1

TableS6.distance_15ca_15ca_same

TableS7.distance_1032sg_1043sg_same

## Code availability

The program is available online on Streamlit cloud:

https://spikescape.streamlitapp.com/

or on our local server:

http://129.49.83.166:8501/

One can also download it directly from the public github repository and use it locally:

https://github.com/stepdasha/spike

The program is written in the Python language^16^ with the use of libraries biotite.v0.34.1,^17^ streamlit.v1.11.1, pandas.v1.4.2^18^ and matplotlib.v3.5.1.^19^

## Acknowledgements

This work was supported by the Research Corporation for Science Advancement (COVID Initiative grant #27350). We gratefully acknowledge support from Henry and Marsha Laufer.

