## Supplemental Lists for "SpikeScape: A Tool for Analyzing Structural Diversity in Experimental Structures of the SARS-CoV-2 Spike Glycoprotein"

### Supplementary information.

#### List S1. Structures removed from all analysis.

There are 66 structures with 1126 not being CYS or 57 not being a PRO in at least one chain:: '6ZB4', '6ZB5', '6ZOZ', '6ZP5', '7CN4', '7EDF', '7EDG', '7EDH', '7EDI', '7EDJ', '7LWQ', '7N9B', '7N9C', '7NS6', '7Q6E', '7Q9F', '7Q9G', '7Q9J', '7Q9M', '7QO7', '7QTI', '7TF1', '7TF2', '7TF4', '7TF5', '7TGW', '7TPL', '7V76', '7V77', '7V7H', '7V7I', '7V7J', '7V7N', '7V7O', '7V7P', '7V7Q', '7V7R', '7V7S', '7V7T', '7V7U', '7V7V', '7V7Z', '7V88', '7V89', '7V8A', '7V8C', '7VRW', '7VXA', '7VXB', '7WE7', '7WE8', '7WE9', '7WEA', '7WEB', '7WEC', '7WG9', '7WJZ', '7WP9', '7WPA', '7WPE', '7XNQ', '7ZR7', '7ZR9', '7ZRC', '8DZH', '8DZI'. Removed those from analysis.

#### List S2. Structures removed from the example 1.

There is 1 chain with missing at least one of the requested 43.N or 565.O atoms: '7CWL'.

#### List S3. Structures removed from the example 2.

There are 45 chains (in 29 structures) with missing at least one of the requested 987C$\alpha$ or 413C$\alpha$ atoms: '6XS6', '6XS6', '6XS6', '6ZOW', '7FB3', '7FB4', '7FB4', '7L3N', '7LAA', '7LAA', '7LSS', '7LSS', '7MTC', '7MTD', '7MTD', '7MTD', '7MTE', '7N9E', '7SXX', '7SXX', '7SXZ', '7SY1', '7SY3', '7SY3', '7SY7', '7T9J', '7T9J', '7T9K', '7TEX', '7TEX', '7TEY', '7TEY', '7TF0', '7TF0', '7TF3', '7TF3', '7THK', '7WLZ', '7WVO', '8DLJ', '8DLM', '8DLM', '8DLX', '8DLZ', '8DLZ'.

#### List S4. Structures with K986P in example 3.

There are 422 structures with K986P: '6VSB', '6VXX', '6VYB', '6WPS', '6WPT', '6X29', '6X2A', '6X2B', '6X2C', '6X6P', '6X79', '6XCM', '6XCN', '6XEY', '6XF5', '6XF6', '6XKL', '6XLU', '6XM0', '6XM3', '6XM4', '6XM5', '6XS6', '6Z43', '6Z97', '6ZDH', '6ZGE', '6ZGG', '6ZGI', '6ZHD', '6ZOW', '6ZOY', '6ZP1', '6ZP7', '6ZXN', '7A25', '7A29', '7A94', '7AKD', '7B18', '7BNM', '7BNN', '7BYR', '7C2L', '7CAB', '7CAC', '7CAI', '7CAK', '7CHH',

'7CT5', '7CYP', '7CZP', '7CZQ', '7CZR', '7CZS', '7CZT', '7CZU', '7CZV', '7CZW', '7CZX', '7CZY', '7CZZ', '7D00', '7D03', '7D0B', '7D0C', '7D0D', '7DDD', '7DF3', '7DF4', '7DK4', '7DWY', '7DWZ', '7DX0', '7DX1', '7DX2', '7DX3', '7DX5', '7DX6', '7DX7', '7DX8', '7DX9', '7DZW', '7DZX', '7DZY', '7E3K', '7E3L', '7E5R', '7E5S', '7E8C', '7E9N', '7E9O', '7E9Q', '7EAZ', '7EB0', '7EB3', '7EB4', '7EB5', '7EH5', '7EJ4', '7EJ5', '7ENF', '7EPX', '7FAE', '7FAF', '7FB0', '7FB1', '7FB3', '7FB4', '7FCD', '7FCE', '7FET', '7FJN', '7FJO', '7JJI', '7JV4', '7JV6', '7JVC', '7JWB', '7JWY', '7JZL', '7JZN', '7K43', '7K4N', '7K8S', '7K8T', '7K8U', '7K8V', '7K8W', '7K8X', '7K8Z', '7K90', '7K9H', '7K9J', '7KJ2', '7KJ3', '7KJ4', '7KJ5', '7KKK', '7KKL', '7KML', '7KMS', '7KMZ', '7KNB', '7KNE', '7KNH', '7KNI', '7KQB', '7KQE', '7KSG', '7L02', '7L06', '7L09', '7L2D', '7L2E', '7L2F', '7L3N', '7L56', '7L7K', '7LAA', '7LAB', '7LCN', '7LD1', '7LJR', '7LQV', '7LRT', '7LS9', '7LSS', '7LXY', '7LXZ', '7LY2', '7M0J', '7M6E', '7M6F', '7M6G', '7M6H', '7M6I', '7MJG', '7MJH', '7MJJ', '7MJK', '7MJM', '7MKL', '7MM0', '7MTC', '7MTD', '7MTE', '7MW2', '7MW3', '7MW4', '7MW5', '7MW6', '7MY2', '7MY3', '7N0G', '7N0H', '7N5H', '7N8H', '7N9E', '7N9T', '7ND3', '7ND4', '7ND5', '7ND7', '7ND8', '7ND9', '7NDA', '7NT9', '7NTA', '7NTC', '7NY5', '7OAN', '7P40', '7P77', '7P78', '7P79', '7P7B', '7Q1Z', '7QDG', '7R13', '7R14', '7R16', '7R17', '7R18', '7R19', '7R1A', '7R1B', '7R40', '7R4I', '7R4Q', '7R4R', '7R8M', '7R8N', '7R8O', '7RA8', '7RBV', '7RKV', '7RU1', '7RU2', '7RU3', '7RU5', '7RW2', '7S0C', '7S0D', '7S6I', '7S6J', '7S6K', '7S6L', '7SC1', '7SN3', '7SOB', '7SOE', '7SXR', '7SXS', '7SXT', '7SXU', '7SXV', '7SXW', '7SXX', '7SXZ', '7SY1', '7SY3', '7SY5', '7SY7', '7T3M', '7T9J', '7T9K', '7TAT', '7TB4', '7TB8', '7TCA', '7TCC', '7TEX', '7TEY', '7TF0', '7TF3', '7TGX', '7TGY', '7THK', '7THT', '7TLA', '7TLB', '7TLC', '7TLD', '7TPR', '7U0P', '7UAP', '7UAR', '7UHC', '7V20', '7V23', '7V26', '7V2A', '7V78', '7V79', '7V7A', '7V7D', '7V7E', '7V7F', '7V7G', '7V81', '7V82', '7V83', '7V85', '7V86', '7VNC', '7VND', '7VNE', '7VQ0', '7VX1', '7VX9', '7VXC', '7VXD', '7VXE', '7VXF', '7VXI', '7VXK', '7VXM', '7W92', '7W94', '7W98', '7W99', '7W9B', '7W9C', '7W9E', '7WCD', '7WCZ', '7WD0', '7WD7', '7WD9', '7WDF', '7WEV', '7WHI', '7WHJ', '7WHK', '7WJY', '7WK2', '7WK3', '7WK4', '7WK5', '7WK9', '7WKA', '7WLY', '7WLZ', '7WO5', '7WOA', '7WOB', '7WPD', '7WPF', '7WS0', '7WS1', '7WS3', '7WS4', '7WS5', '7WS8', '7WS9', '7WVN', '7WVO', '7WVP', '7WWL', '7WWM', '7WZ1', '7WZ2', '7X08', '7X6A', '7XCH', '7XCO', '7XIC', '7XID', '7XIW', '7XIX', '7XIY', '7XNR', '7XNS', '7XO4', '7XO5', '7XO7', '7XO8', '7XOA', '7XOB', '7XOD', '7Y9S', '7Y9Z', '7YA0', '7YEG', '7YQT', '7YQU', '7YQV', '7YQW', '7YQX',

'7YQY', '7YQZ', '7YR1', '7YR2', '7YR3', '7Z3Z', '7Z6V', '7Z7X', '7Z85', '7Z86', '7Z9Q', '7ZCE', '8CXN', '8CXQ', '8CY6', '8CY7', '8CY9', '8CYA', '8CYB', '8CYC', '8CYD', '8DLI', '8DLJ', '8DLL', '8DLM', '8DLO', '8DLP', '8DLT', '8DLU', '8DLW', '8DLX', '8DLZ'

#### List S5. Structures removed from the example 3.

There are 45 chains (in 29 structures) with missing at least one of the requested 987.C$\alpha$ or 413.C$\alpha$ in K986P structures: '6XS6', '6XS6', '6XS6', '6ZOW', '7FB3', '7FB4', '7FB4', '7L3N', '7LAA', '7LAA', '7LSS', '7LSS', '7MTC', '7MTD', '7MTD', '7MTD', '7MTE', '7N9E', '7SXX', '7SXX', '7SXZ', '7SY1', '7SY3', '7SY3', '7SY7', '7T9J', '7T9J', '7T9K', '7TEX', '7TEX', '7TEY', '7TEY', '7TF0', '7TF0', '7TF3', '7TF3', '7THK', '7WLZ', '7WVO', '8DLJ', '8DLM', '8DLM', '8DLX', '8DLZ', '8DLZ'. 420 structures we used for distance distribution calculation.

#### List S6. Structures with K986 in example 3.

There are 97 structures with K986: '6XR8', '6ZOX', '6ZP0', '6ZWV', '7A4N', '7AD1', '7CWL', '7CWM', '7CWN', '7CWS', '7CWT', '7CWU', '7E7B', '7E7D', '7KDG', '7KDH', '7KDI', '7KDJ', '7KDK', '7KDL', '7KE4', '7KE6', '7KE7', '7KE8', '7KE9', '7KEA', '7KEB', '7KEC', '7KRQ', '7KRR', '7KRS', '7LWI', '7LWJ', '7LWK', '7LWL', '7LWM', '7LWN', '7LWO', '7LWP', '7LWS', '7LWT', '7LWU', '7LWV', '7LWW', '7LYK', '7LYL', '7LYM', '7LYN', '7LYO', '7LYQ', '7N1Q', '7N1T', '7N1U', '7N1V', '7N1W', '7N1X', '7OD3', '7ODL', '7QUR', '7QUS', '7SBK', '7SBL', '7SBO', '7SBP', '7SBQ', '7SBR', '7SBS', '7SBT', '7SO9', '7TEI', '7TM0', '7TNW', '7TO4', '7TOU', '7TOV', '7TOX', '7TOY', '7TP0', '7TP1', '7TP2', '7TP7', '7TP8', '7TP9', '7TPA', '7TPC', '7TPE', '7TPF', '7TPH', '7UB0', '7UB5', '7UB6', '7UPW', '7UPY', '7WG7', '7WGB', '7XU1', '8CSA'.

#### List S7. Structures with missing 15C$\alpha$ atom from the example 4.

There are 880 chains (in 298 structures) with missing 15C$\alpha$ atom: '6VSB', '6VSB', '6VSB', '6VXX', '6VXX', '6VXX', '6VYB', '6VYB', '6VYB', '6WPS', '6WPS', '6WPS', '6WPT', '6WPT', '6WPT', '6X29', '6X29', '6X29', '6X2A', '6X2A', '6X2A', '6X2B', '6X2B', '6X2B', '6X2C', '6X2C', '6X2C', '6X6P',

'6X6P', '6X6P', '6X79', '6X79', '6X79', '6XCM', '6XCM', '6XCM', '6XCN', '6XCN', '6XCN', '6XEY', '6XEY', '6XEY', '6XF5', '6XF5', '6XF5', '6XF6', '6XF6', '6XF6', '6XKL', '6XKL', '6XKL', '6XLU', '6XLU', '6XLU', '6XM0', '6XM0', '6XM0', '6XM3', '6XM3', '6XM3', '6XM4', '6XM4', '6XM4', '6XM5', '6XM5', '6XM5', '6XS6', '6XS6', '6XS6', '6Z43', '6Z43', '6Z43', '6Z97', '6Z97', '6Z97', '6ZDH', '6ZDH', '6ZDH', '6ZHD', '6ZHD', '6ZHD', '6ZOW', '6ZOW', '6ZOW', '6ZOX', '6ZOX', '6ZOX', '6ZOY', '6ZOY', '6ZOY', '6ZP0', '6ZP0', '6ZP0', '6ZP1', '6ZP1', '6ZP1', '6ZP7', '6ZP7', '6ZP7', '6ZWV', '6ZWV', '6ZWV', '7A4N', '7A4N', '7A4N', '7AD1', '7AD1', '7AD1', '7AKD', '7AKD', '7AKD', '7B18', '7BYR', '7BYR', '7BYR', '7CAB', '7CAB', '7CAB', '7CAC', '7CAC', '7CAC', '7CAI', '7CAI', '7CAI', '7CAK', '7CAK', '7CAK', '7CHH', '7CHH', '7CHH', '7CT5', '7CT5', '7CT5', '7CYP', '7CYP', '7CYP', '7CZP', '7CZP', '7CZP', '7CZQ', '7CZQ', '7CZQ', '7CZR', '7CZR', '7CZR', '7CZS', '7CZS', '7CZS', '7CZT', '7CZT', '7CZT', '7CZU', '7CZU', '7CZU', '7CZV', '7CZV', '7CZV', '7CZW', '7CZW', '7CZW', '7CZX', '7CZX', '7CZX', '7CZY', '7CZY', '7CZY', '7CZZ', '7CZZ', '7CZZ', '7D00', '7D00', '7D00', '7D03', '7D03', '7D03', '7D0B', '7D0B', '7D0B', '7D0C', '7D0C', '7D0C', '7D0D', '7D0D', '7D0D', '7DF4', '7DWZ', '7DWZ', '7DWZ', '7DX0', '7DX0', '7DX0', '7DX1', '7DX1', '7DX1', '7DX2', '7DX2', '7DX2', '7DX3', '7DX3', '7DX3', '7DX5', '7DX5', '7DX5', '7DX6', '7DX6', '7DX6', '7DX7', '7DX7', '7DX7', '7DX8', '7DX8', '7DX8', '7DX9', '7DX9', '7DX9', '7DZW', '7DZW', '7DZW', '7DZX', '7DZX', '7DZX', '7DZY', '7DZY', '7DZY', '7E3K', '7E3K', '7E3K', '7E3L', '7E3L', '7E3L', '7E9N', '7E9N', '7E9N', '7E9O', '7E9O', '7E9O', '7E9Q', '7E9Q', '7E9Q', '7EAZ', '7EAZ', '7EAZ', '7EB0', '7EB0', '7EB0', '7EB3', '7EB3', '7EB3', '7EB4', '7EB4', '7EB4', '7EB5', '7EB5', '7EB5', '7EH5', '7EH5', '7EH5', '7EJ4', '7EJ4', '7EJ4', '7EJ5', '7EJ5', '7EJ5', '7ENF', '7ENF', '7ENF', '7EPX', '7EPX', '7EPX', '7FAE', '7FAE', '7FAE', '7FAF', '7FAF', '7FAF', '7FB0', '7FB0', '7FB0', '7FB1', '7FB1', '7FB1', '7FB3', '7FB3', '7FB3', '7FB4', '7FB4', '7FB4', '7FET', '7FET', '7FET', '7FJN', '7FJN', '7FJN', '7FJO', '7FJO', '7FJO', '7JV4', '7JV4', '7JV4', '7JV6', '7JV6', '7JV6', '7JVC', '7JVC', '7JVC', '7JWB', '7JWB', '7JWB', '7JWY', '7JWY', '7JWY', '7JZL', '7JZL', '7JZL', '7JZN', '7JZN', '7JZN', '7K4N', '7K4N', '7K4N', '7K8S', '7K8S', '7K8S', '7K8T', '7K8T', '7K8T', '7K8U', '7K8U', '7K8U', '7K8V', '7K8V', '7K8V', '7K8W', '7K8W', '7K8W', '7K8X', '7K8X', '7K8X', '7K8Z', '7K8Z', '7K8Z', '7K90', '7K90', '7K90', '7K9H', '7K9H', '7K9H', '7K9J', '7K9J', '7K9J', '7KDG', '7KDG', '7KDG', '7KDH', '7KDH', '7KDH', '7KDI', '7KDI', '7KDI', '7KDJ', '7KDJ', '7KDJ', '7KDK', '7KDK', '7KDK', '7KDL', '7KDL', '7KDL', '7KE4', '7KE4', '7KE4', '7KE6',

'7KE6', '7KE6', '7KE7', '7KE7', '7KE7', '7KE8', '7KE8', '7KE8', '7KE9', '7KE9', '7KE9', '7KEA', '7KEA', '7KEA', '7KEB', '7KEB', '7KEB', '7KEC', '7KEC', '7KEC', '7KJ2', '7KJ2', '7KJ2', '7KJ3', '7KJ3', '7KJ3', '7KJ4', '7KJ4', '7KJ4', '7KJ5', '7KJ5', '7KJ5', '7KKK', '7KKK', '7KKK', '7KKL', '7KKL', '7KKL', '7KMS', '7KMS', '7KMS', '7KMZ', '7KMZ', '7KMZ', '7KNB', '7KNB', '7KNB', '7KNE', '7KNE', '7KNE', '7KNH', '7KNH', '7KNH', '7KNI', '7KNI', '7KNI', '7KSG', '7L02', '7L02', '7L02', '7L06', '7L06', '7L06', '7L09', '7L09', '7L09', '7L3N', '7L3N', '7L3N', '7L56', '7L56', '7L56', '7L7K', '7L7K', '7L7K', '7LAA', '7LAA', '7LAA', '7LAB', '7LAB', '7LAB', '7LCN', '7LCN', '7LCN', '7LD1', '7LD1', '7LD1', '7LJR', '7LJR', '7LJR', '7LSS', '7LSS', '7LSS', '7LWI', '7LWI', '7LWI', '7LWJ', '7LWJ', '7LWJ', '7LWK', '7LWK', '7LWK', '7LWL', '7LWL', '7LWL', '7LWM', '7LWM', '7LWM', '7LWN', '7LWN', '7LWN', '7LWO', '7LWO', '7LWO', '7LWP', '7LWP', '7LWP', '7LWS', '7LWS', '7LWS', '7LWT', '7LWT', '7LWT', '7LWU', '7LWU', '7LWU', '7LWV', '7LWV', '7LWV', '7LWW', '7LWW', '7LWW', '7LYK', '7LYK', '7LYK', '7LYL', '7LYL', '7LYL', '7LYM', '7LYM', '7LYM', '7LYN', '7LYN', '7LYN', '7LYO', '7LYO', '7LYO', '7LYQ', '7LYQ', '7LYQ', '7M0J', '7M0J', '7M0J', '7M6F', '7M6F', '7M6F', '7M6G', '7M6G', '7M6G', '7M6H', '7M6H', '7M6H', '7M6I', '7M6I', '7M6I', '7MKL', '7MKL', '7MKL', '7MTC', '7MTC', '7MTC', '7MTD', '7MTD', '7MTD', '7MTE', '7MTE', '7MTE', '7MW2', '7MW2', '7MW2', '7MW3', '7MW3', '7MW3', '7MW4', '7MW4', '7MW4', '7MY2', '7MY2', '7MY2', '7MY3', '7MY3', '7MY3', '7N5H', '7N5H', '7N5H', '7N8H', '7N8H', '7N8H', '7N9E', '7N9E', '7N9E', '7N9T', '7N9T', '7N9T', '7ND3', '7ND3', '7ND3', '7ND4', '7ND4', '7ND4', '7ND5', '7ND5', '7ND5', '7ND7', '7ND7', '7ND7', '7ND8', '7ND8', '7ND8', '7ND9', '7ND9', '7ND9', '7NDA', '7NDA', '7NDA', '7NY5', '7NY5', '7NY5', '7OD3', '7OD3', '7OD3', '7ODL', '7ODL', '7ODL', '7P40', '7P40', '7P40', '7P77', '7P77', '7P77', '7P78', '7P78', '7P78', '7P79', '7P79', '7P79', '7Q1Z', '7Q1Z', '7Q1Z', '7R13', '7R13', '7R13', '7R14', '7R14', '7R14', '7R16', '7R16', '7R16', '7R17', '7R17', '7R17', '7R1A', '7R1A', '7R1A', '7R8M', '7R8M', '7R8M', '7RA8', '7RA8', '7RA8', '7RKV', '7RKV', '7RKV', '7RU3', '7S0C', '7S0C', '7S0C', '7S6L', '7S6L', '7S6L', '7SC1', '7SC1', '7SC1', '7SN3', '7SN3', '7SN3', '7T3M', '7T3M', '7T3M', '7TAT', '7TAT', '7TAT', '7TEI', '7TEI', '7TEI', '7THK', '7THK', '7THK', '7THT', '7THT', '7THT', '7TLA', '7TLA', '7TLA', '7TLB', '7TLB', '7TLB', '7TLC', '7TLC', '7TLC', '7TLD', '7TLD', '7TLD', '7TOU', '7TOU', '7TOU', '7TOV', '7TOV', '7TOV', '7TOX', '7TOX', '7TOX', '7TOY', '7TOY', '7TOY', '7TP0', '7TP0', '7TP0', '7TP1', '7TP1', '7TP1', '7TP2', '7TP2', '7TP2', '7TP7', '7TP7', '7TP7', '7TP8', '7TP8', '7TP8', '7TP9', '7TP9', '7TP9', '7TPA', '7TPA', '7TPA',

'7TPC', '7TPC', '7TPC', '7TPE', '7TPE', '7TPE', '7TPF', '7TPF', '7TPF', '7TPH', '7TPH', '7TPH', '7UAR', '7UAR', '7UAR', '7UB0', '7UB0', '7UB0', '7UB5', '7UB5', '7UB5', '7UB6', '7UB6', '7UB6', '7UHC', '7UHC', '7UHC', '7V20', '7V20', '7V20', '7V2A', '7V2A', '7V2A', '7WCD', '7WCD', '7WCD', '7WHI', '7WHJ', '7WHK', '7WJY', '7WJY', '7WJY', '7WLY', '7WLY', '7WLY', '7WLZ', '7WLZ', '7WLZ', '7WPD', '7WPD', '7WPD', '7WPF', '7WPF', '7WPF', '7WWL', '7WWL', '7WWL', '7WWM', '7WWM', '7WWM', '7WZ1', '7WZ1', '7WZ1', '7X6A', '7X6A', '7X6A', '7XIC', '7XIC', '7XIC', '7XID', '7XID', '7XID', '7XIW', '7XIW', '7XIW', '7XIX', '7XIX', '7XIX', '7XIY', '7XIY', '7XIY', '7XNR', '7XNR', '7XNR', '7XNS', '7XNS', '7XNS', '7XO4', '7XO4', '7XO4', '7XO5', '7XO5', '7XO5', '7XO7', '7XO7', '7XO7', '7XO8', '7XO8', '7XO8', '7XOA', '7XOA', '7XOA', '7XOB', '7XOB', '7XOB', '7XOD', '7XOD', '7XOD', '7YQT', '7YQT', '7YQT', '7YQU', '7YQU', '7YQU', '7YQV', '7YQV', '7YQV', '7YQW', '7YQW', '7YQW', '7YQX', '7YQX', '7YQX', '7YQY', '7YQY', '7YQY', '7YQZ', '7YQZ', '7YQZ', '7YR1', '7YR1', '7YR1', '7YR2', '7YR2', '7YR2', '7YR3', '7YR3', '7YR3', '7Z6V', '7Z6V', '7Z6V', '7Z7X', '7Z7X', '7Z7X', '7Z85', '7Z85', '7Z85', '7Z86', '7Z86', '7Z86', '7Z9Q', '7Z9Q', '7Z9Q', '7ZCE', '7ZCE', '7ZCE', '8CSA', '8CSA', '8CSA', '8CY7', '8CY7', '8CY7'.

#### List S8. Structures removed from the example 5.

There are 9 chains (in 3 structures) with missing at least one of the requested CYS1043:SG or CYS1032:SG atoms: '7DZW', '7DZW', '7DZW', '7DZX', '7DZX', '7DZX', '7DZY', '7DZY', '7DZY'. 516 structures we used for distance distribution calculation.
